# Language laterality and cognitive skills: does anatomy matter?

**DOI:** 10.1101/2025.09.06.674566

**Authors:** Ieva Andrulyte, Laure Zago, Gael Jobard, Herve Lemaitre, Francesca M. Branzi, Francois Rheault, Laurent Petit, Simon S Keller

**Affiliations:** Department of Pharmacology and Therapeutics, Institute of Systems, Molecular and Integrative Biology, University of Liverpool, Liverpool, UK; Universite de Bordeaux, CNRS, CEA, IMN, UMR 5293, GIN, F-33000 Bordeaux, France; Department of Psychological Sciences, Institute of Population Health, University of Liverpool, Liverpool, UK; Université de Sherbrooke, Department of Computer Science, Sherbrooke, Québec, Canada; IRP OpTeam, CNRS Biologie, France - Université de Sherbrooke, Canada

**Keywords:** language lateralisation, cognitive function, diffusion MRI, white matter, arcuate fasciculus, corpus callosum

## Abstract

Brain anatomy, particularly white matter microstructure, is thought to play a critical role in the relationships between cognitive function and language lateralisation. This study investigates whether white matter microstructural parameters of the arcuate fasciculus and corpus callosum is associated with cognitive performance across distinct language lateralisation groups. Neuroimaging and cognitive data from 279 healthy adults were sourced from the BIL&GIN database. Participants completed a sentence production fMRI task, diffusion MRI, and cognitive tasks assessing verbal, visuospatial, and arithmetic skills. Significant positive associations were observed in strongly atypical individuals between fractional anisotropy in the splenium and working memory (R^2^ = 0.96, pFDR = 0.042) and between FA in the genu and visuospatial attention (R^2^ = 0.92, pFDR = 0.042). ANCOVA revealed that cross-dominant individuals had significantly lower visuospatial attention scores compared to consistently lateralised individuals (pFDR = 0.02). These findings challenge the notion that atypical lateralisation is inherently maladaptive and suggest that white matter pathways may serve as an alternative mechanism for supporting cognitive function in individuals with rightward language dominance. Furthermore, the results highlight the cognitive disadvantages of crossed dominance, implicating disrupted interhemispheric communication as a potential underlying mechanism.

## Introduction

Language lateralisation—the preferential involvement of one hemisphere in language processing—is a fundamental feature of human neurobiology. In most right-handed individuals and a large proportion of left-handers, language functions are predominantly localised in the left hemisphere (Knecht, Dräger, et al., 2000; Mazoyer et al., 2014; Olulade et al., 2020). It has been suggested that left-hemispheric dominance may enhance cognitive efficiency by minimising interhemispheric transfer times and supporting rapid linguistic processing (Gotts et al., 2013). However, lateralisation patterns vary, with some individuals exhibiting bilateral or right-hemispheric dominance, and the mechanisms underlying this variability remain unclear. While left lateralisation has been linked to cognitive advantages in verbal fluency, spatial reasoning, and memory (Gotts et al., 2013; Mellet et al., 2014), other studies report no significant association between lateralisation and cognitive abilities such as linguistic processing, intelligence, or academic achievement (Cai et al., 2013; Knecht, Deppe, et al., 2000). These inconsistencies suggest that language lateralisation is not a direct predictor of cognitive performance and may instead interact with broader neural and structural factors (Bishop, 2013).

Language lateralisation is often described at the whole-brain level, but increasing evidence suggests that lateralisation patterns can vary across different brain regions. Studies examining regional lateralisation have shown that different cortical areas can exhibit distinct lateralisation patterns, which may not align with whole-brain measures. For example, Van Ettinger-Veenstra et al. (2012) found that right-lateralised activation in the dorsolateral prefrontal region was associated with better linguistic abilities compared to left-lateralised activation in the same region, challenging the conventional assumption that left-hemispheric dominance is always beneficial for language function and indicating that language networks may be more flexible than previously assumed (Bradshaw, Bishop, et al., 2017). One manifestation of this variability is crossed language dominance, where language-related activation is lateralised to opposite hemispheres in the frontal and temporal lobes (Seghier, 2008). While people with crossed language dominance are rare in the healthy population and is often considered a disruption of the expected lateralisation pattern (Berl et al., 2014), it remains unclear whether crossed language dominance reflects functional reorganisation, an adaptive mechanism, or a marker of neural inefficiency. Furthermore, how these variations relate to cognitive function and anatomical differences remains an open question.

White matter microstructure plays a key role in shaping both language lateralisation and cognitive function (Andrulyte et al., 2024; Raghavan et al., 2020), with two tracts—the arcuate fasciculus and corpus callosum—central to these processes. The arcuate fasciculus, a frontotemporal association pathway, facilitates intrahemispheric communication between core language regions, while the CC, the brain’s largest commissural tract, integrates linguistic and cognitive information across hemispheres, balancing excitatory and inhibitory interactions (Friederici et al., 2011; Hickok & Poeppel, 2007; Perani et al., 2011; Ribeiro et al., 2024). Both tracts have been implicated in language lateralisation, yet their precise role remains debated. Some studies suggest that higher arcuate fasciculus fractional anisotropy (FA) supports left-hemispheric dominance (Ocklenburg et al., 2013), while others find no clear association (Gerrits et al., 2021; Verhelst et al., 2021). Similarly, the corpus callosum role is disputed, with some studies linking greater microstructural measures to stronger functional lateralisation (Hinkley et al., 2016; Josse et al., 2008; Karolis et al., 2019), while others suggest it facilitates interhemispheric communication, potentially related to reduced lateralisation (Andrulyte et al., 2024; Häberling et al., 2011; Zhu et al., 2025).

Both the arcuate fasciculus and corpus callosum are also associated with cognitive abilities, particularly in language processing, general cognition, and attention (Catani et al., 2007; Forkel et al., 2022; Niogi et al., 2008; Villar-Rodríguez, Marin-Marin, et al., 2024). Greater arcuate fasciculus microstructural measures have been linked to improved verbal fluency and learning (Lebel & Beaulieu, 2009; López-Barroso et al., 2013; Salvan et al., 2017), while corpus callosum microstructural properties have been associated with improved linguistic and executive function (Bartha-Doering et al., 2021; Danielsen et al., 2020; Pliatsikas et al., 2015; Ribeiro et al., 2024). Conversely, damage to these tracts has been linked to cognitive impairments—lesions in arcuate fasciculus disrupt fluency, repetition, and linguistic processing (Forkel et al., 2014; Hillis et al., 2018; Hosomi et al., 2009; Paldino et al., 2016; Vavassori et al., 2023), while corpus callosum lesions impair interhemispheric transfer and higher-order cognitive abilities (Fabri et al., 2025; Sidaros et al., 2008). Despite these established links, no studies have examined how white matter structure influences both cognition and language lateralisation simultaneously, particularly in individuals with atypical or crossed dominance.

It is unclear whether white matter microstructure differentially influences cognitive performance across individuals with typical and atypical lateralisation, or whether anatomical variations in the arcuate fasciculus and corpus callosum contribute to crossed language dominance. The cognitive profiles of individuals with crossed dominance also remain poorly characterised, particularly in relation to white matter microstructural parameters and interhemispheric connectivity. This study addresses these gaps by investigating the interplay between white matter anatomy, cognition, and language lateralisation in a large sample of healthy adults. Specifically, we examine whether (1) white matter microstructure mediates the association between cognitive function and different lateralisation patterns, (2) FA in the corpus callosum and arcuate fasciculus relates to cognitive abilities across different lateralisation groups, and (3) individuals with consistent lateralisation differ from those with crossed dominance in cognitive performance and white matter microstructural parameters.

## Methods

### Participants

Structural and functional neuroimaging data for this study were sourced from the BIL&GIN database, a multimodal imaging resource specifically designed to investigate the structural and functional neural correlates of brain lateralisation (Mazoyer et al., 2016). The database includes cognitive, diffusion MRI, and functional MRI data from healthy adults with no history of brain abnormalities (Figure 1). Ethical approval for the database was granted by the Basse-Normandie local Ethics Committee. For our study, we included participants with complete data across sentence production task fMRI, diffusion MRI, and cognitive tasks. This selection resulted in a final sample of 279 participants (137 females; age range: 19– 57 years; mean age: 25.7 years; SD = 6.5). The cohort comprised a balanced distribution of handedness, with 133 right-handed and 146 left-handed individuals (χ^2^ = 0.6, p = 0.43).

**Figure 1.**
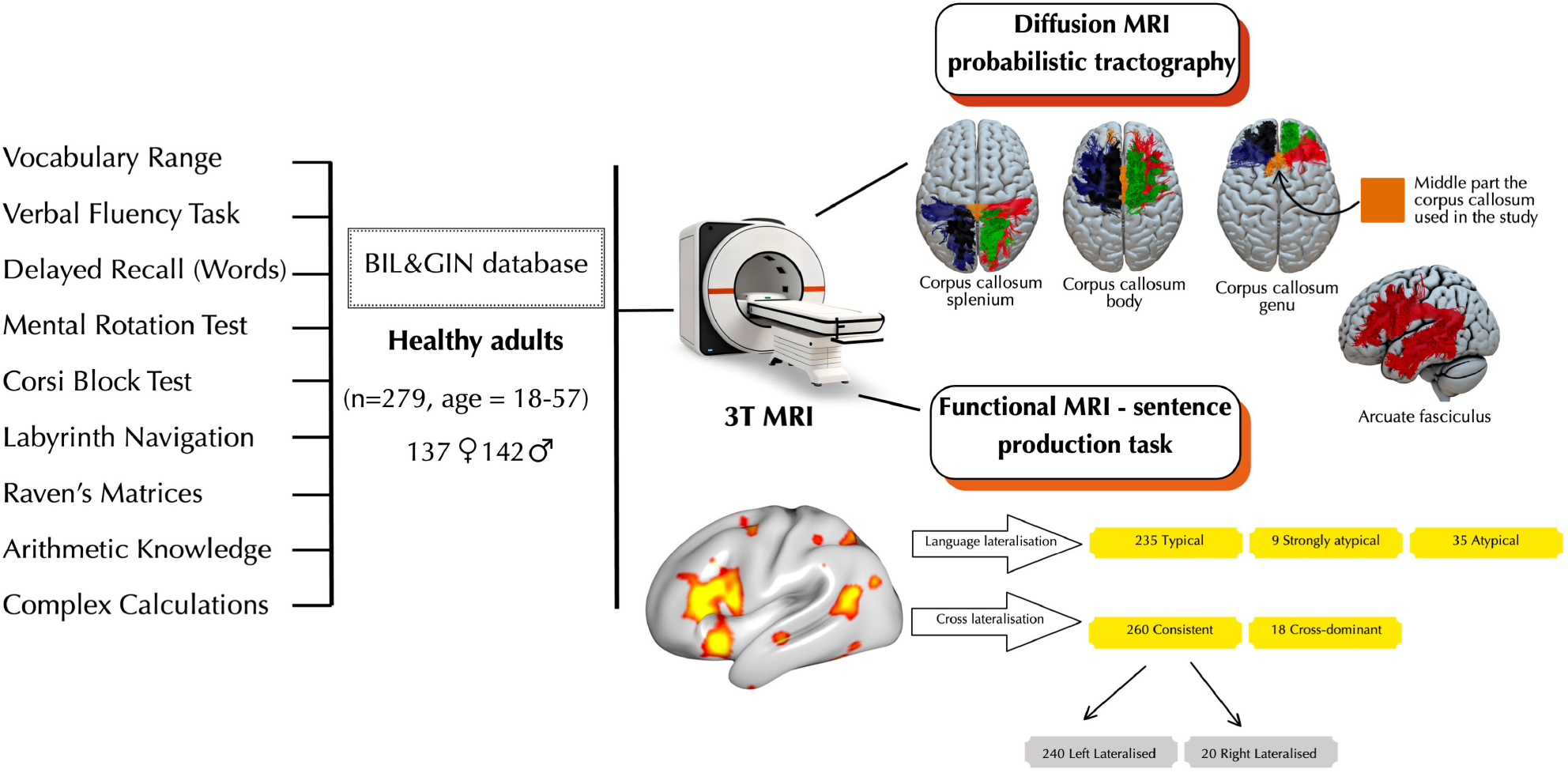
Overview of data and modalities used. This study utilised the BIL&GIN database, including 279 healthy adults (137 females, 142 males, aged 18–57). Cognitive tasks assessed language processing, working memory, spatial orientation, and mathematical thinking. Diffusion MRI probabilistic tractography was performed on the arcuate fasciculus and corpus callosum (body, genu, and splenium), with the AF and CC reconstructions presented in this figure derived from a single participant in the BIL&GIN dataset. These reconstructions serve as an example, recognising that AF and CC structures vary across individuals. Functional MRI (sentence production task) determined language lateralisation, categorising participants as Typical (n=235), Atypical (n=35), or Strongly Atypical (n=9). Cross-lateralisation analysis identified Consistently Lateralised (n=260) and Cross-Dominant (n=18) individuals, with 240 being left-lateralised and 20 right-lateralised.

### MRI acquisition and processing

Neuroimaging data were acquired using a Philips Achieva 3-Tesla MRI scanner, optimised to provide high-quality structural and functional brain images. Structural imaging included a high-resolution 3D T1-weighted sequence (TR = 20 ms, TE = 4.6 ms, flip angle = 10°, inversion time = 800 ms, turbo field echo factor = 65, SENSE factor = 2, matrix size = 256 × 256 × 180, and isotropic voxel size = 1 mm^3^).

Functional MRI data were obtained using T2*-weighted echo-planar imaging (EPI) with parameters tailored for capturing brain activity: TR = 2 s, TE = 35 ms, flip angle = 80°, and 31 axial slices, each with an isotropic voxel size of 3.75 mm^3^. During the fMRI session, participants completed a sentence production task (Mazoyer et al., 2016), which involved three runs of 192 volumes. To ensure optimal data quality, the first four volumes of each run were discarded, allowing the MR signal to stabilise.

Diffusion-weighted imaging (DWI) was conducted using a spin-echo EPI sequence comprising a b0 image (b = 0 s/mm^2^) and 21 distinct diffusion-weighted directions (b = 1000 s/mm^2^), each acquired twice with reversed phase-encoding polarities, resulting in a total of 84 images. Imaging covered 70 axial slices aligned to the anterior commissure–posterior commissure (AC–PC) plane, spanning the brain from the cerebellum to the vertex. Acquisition parameters included TR = 8500 ms, TE = 81 ms, flip angle = 90°, SENSE factor = 2.5, field of view = 224 mm, matrix size = 112 × 112, and isotropic voxel size of 2 mm^3^. The protocol was repeated to improve signal-to-noise ratio, with a total scan time of approximately 15 minutes and 30 seconds.

### fMRI data processing

Task-based fMRI data were processed using Statistical Parametric Mapping (SPM5) software and custom MATLAB scripts. Functional data from the three slow fMRI runs were corrected for slice timing and motion. To account for motion artefacts, six movement parameters (three translations and three rotations) were regressed out from the T2*-weighted EPI time series. The corrected functional images were then co-registered to participants’ structural T1-weighted images. Spatial normalisation was performed by combining the co-registration matrix with the transformation matrix from structural images to standard stereotaxic space, resulting in normalised T2*-EPI scans with a voxel resolution of 2 × 2 × 2 mm^3^. Normalisation employed trilinear interpolation to ensure accurate warping. A high-pass filter with a 159-second cut-off was applied to remove low-frequency noise (Joliot et al., 2016).

### fMRI language paradigm

During the sentence production task (PROD_SENT_), participants were shown a line drawing for 1 second on a black screen, sourced from the French comic series Le Petit Nicolas. Participants were instructed to covertly generate a structured sentence starting with a subject and complement (e.g., “The little Nicolas…”), followed by a verb and an adverbial phrase of place or manner (e.g., “in the street…”, “with happiness…”). In the reference word-list production task (PROD_LIST_), scrambled versions of the drawings were presented as stimuli. Participants were required to covertly recite ordered lists, such as the months of the year, and press a response pad upon completion. Both tasks lasted 8-14 seconds per trial. Following both tasks, participants performed a low-level visuo-motor baseline task, detecting the transformation of a central fixation cross into a square and pressing a button upon noticing the change. This baseline phase was designed to last at least half of the total trial duration, controlling for motor responses and maintaining attention. Each trial lasted 18 seconds, including the stimulus presentation, response period, and baseline task. A 12-second fixation cross was presented before and after the first and last trials to ensure focus. Further details are available in Mazoyer et al., (2014).

### Language production lateralisation

Language laterality indices (LIs) for this study were calculated using the LI toolbox (Wilke & Schmithorst, 2006) applied to the individual PROD_SENT_ minus PROD_LIST_ t-maps. For more information please refer to (Mazoyer et al., 2014). Final LIs, ranging from −100 (indicating complete right-lateralisation) to +100 (indicating complete left-lateralisation), were classified into three categories: individuals with LI values above 18 were classified as “typically” lateralised, those between −50 and 18 as “atypically” lateralised, and those below −50 as “strongly atypical,” following criteria established in Mazoyer et al. (2014).

To study the phenomenon of cross laterality, we utilised the LIs for frontal and temporal regions of interest (ROIs), as defined by the SENSAAS language atlas (Labache et al., 2019), which was created using BIL&GIN dataset, based on PROD_SENT_ minus PROD_LIST_ t-maps. For the frontal region, the LI for the pars triangularis of the inferior frontal gyrus (F3t) was utilised. Although the pars opercularis is known to play a role in language production, it was not included in our analysis as it was not identified as a key region for sentence production in the SENSAAS atlas. For the temporal region, the LIs for superior temporal sulcus 3 (STS3) and 4 (STS4) were combined by averaging their values. For categorisation, we applied thresholds of 10, where values greater than the positive threshold indicated left-lateralised language dominance, values below the negative threshold (-10) indicated right-lateralised dominance, and values between these thresholds were categorised as bilateral (Vernooij et al., 2007). Participants were categorised as crossed-dominant if their frontal and temporal regions were lateralised to opposite hemispheres (e.g., frontal region left-lateralised and temporal region right-lateralised, or vice versa) (Seghier, 2008; Tailby et al., 2017). Those with the same lateralisation in both regions, including cases where one or both regions were classified as bilateral, were categorised as consistent lateralisation, as these did not involve switching between hemispheres.

### Diffusion MRI analysis

Probabilistic tractography was conducted using TractoFlow (Theaud et al., 2020), an automated diffusion MRI processing pipeline. TractoFlow integrates two tracking algorithms—classical local tracking and Particle Filter Tracking (PFT) (Girard et al., 2014)—which were used together to generate whole-brain tractograms. Fibre orientation distribution functions (fODFs) were estimated using constrained spherical deconvolution with a maximum spherical harmonic order of 6 (Descoteaux et al., 2009). The resulting tractograms were normalised to MNI space through linear and non-linear transformations and the ExTractorFlow pipeline was used to eliminate anatomically implausible tracts by applying Boolean rules that isolate streamlines meeting specific anatomical criteria (Petit et al., 2023). For more details on diffusion MRI preprocessing please refer to Andrulyte, Zago, et al. (2025).

The arcuate fasciculus and corpus callosum white matter tracts were reconstructed using anatomical definitions from the JHU atlas (Oishi et al., 2009). The corpus callosum was reconstructed by extracting homotopic callosal fibres connecting corresponding gyri of the left and right hemispheres using the JHU ICM atlas (Petit et al., 2023). Recognising that different corpus callosum subregions contribute uniquely to language lateralisation (Andrulyte et al., 2024; Karpychev et al., 2022), the corpus callosum was segmented into three divisions—genu (GCC), body (BCC), and splenium (SCC)—following an anterior-to-posterior axis using JHU atlas. Within each of these three regions, the tracts were further divided into five equally spaced subdivisions along their longitudinal (horizontal) axis. This process involved computing a centroid for each tract to serve as a reference axis, followed by dividing the tracts into five parts along this axis. For our analyses, we exclusively focused on the third (central) subdivision of the genu, body, and splenium of the corpus callosum. The third segment aligns closely with the mid-sagittal plane, which is widely used in MRI studies due to its ability to summarise the structural properties of all callosal fibres (Xiong et al., 2024). Additionally, using the central subdivision mitigates partial volume effects, which can arise when streamlines from adjacent regions are mixed (Kim et al., 2008; Stadler et al., 2023).

The FA of the left and right arcuate fasciculus, as well as the genu (GCC), body (BCC), and splenium (SCC) of the corpus callosum, were calculated using the FSL software package (Jenkinson et al., 2012). Binary masks were generated for each tract segment to delineate voxels corresponding to the streamlines of interest. These masks were then applied to subject-specific FA maps to extract the mean FA values for the selected tracts. While FA is known to be influenced by factors such as crossing fibres and partial volume effects (Jeurissen et al., 2013), it remains the most extensively used diffusion metric in studies investigating language lateralisation (Andrulyte, Demirkan, et al., 2025), thus enabling a better cross-study comparison.

### Cognitive battery

The BIL&GIN cognitive battery includes a diverse set of tasks designed to evaluate various cognitive and language-related abilities. For this study, we selected 9 tasks, encompassing verbal, visuospatial, and arithmetic abilities, as shown in Figure 1. All selected tasks are inherently tied to language skills, either through their reliance on linguistic processing or their connection to cognitive functions mediated by language. Further methodological details are available in previous studies (Labache et al., 2020; Mazoyer et al., 2016).

Verbal skills were assessed using three tasks, capturing dimensions of verbal memory and semantic fluency: (1) vocabulary range test looked at lexical knowledge and semantic richness through a synonym-finding task; (2) verbal fluency measured participants’ ability to generate verbs for given nouns, reflecting semantic retrieval efficiency; and (3) delayed recall words task assessed episodic memory.

Visuospatial skills were assessed using four tasks: (1) mental rotation test measured spatial manipulation by comparing 3D objects (Vandenberg & Kuse, 1978); (2) Corsi block test evaluated visuospatial short-term memory by requiring participants to replicate sequences of block taps (Della Sala et al., 1999); (3) labyrinth navigation assessed spatial problem-solving and topographic orientation; and (4) Raven’s matrices measured abstract reasoning and pattern recognition through completion of geometric matrices (Raven, 1956). Arithmetic knowledge test involved solving basic arithmetic problems, such as multiplication, testing numerical recall, while complex calculations test measured higher-order mathematical reasoning through multi-step problem-solving.

### Statistical analyses

All statistical analyses were conducted in R (version 4.4.1). Principal component analysis (PCA) was conducted to reduce the dimensionality of the cognitive test data while retaining the variance across variables. PCA identifies principal components (PCs)—uncorrelated variables that summarise patterns in the original data. To ensure comparability across cognitive test scores measured on different scales, the data were standardised prior to analysis. PCA was performed using the prcomp() function, which employs orthogonal decomposition to compute eigenvalues and eigenvectors, indicating the amount of variance explained by each component. The number of components to retain was determined using the standard criterion of selecting those with eigenvalues greater than 1, ensuring that only meaningful components were included. A scree plot was generated to visually assess the eigenvalues and confirm the retained components.

To investigate the relationships between cognitive performance, lateralisation, and white matter microstructure, we employed both ANCOVA and multiple regression to address distinct but complementary research questions. ANCOVA was used to assess group-level differences in cognitive PC scores while controlling for individual variability in FA, whereas multiple regression was applied to examine continuous associations between FA and cognitive performance within each lateralisation subgroup. The combined use of these methods ensured appropriate separation of between-group and within-group variance components, preventing the conflation of categorical effects (group differences in cognition) with continuous associations (individual differences in FA) (Cohen, 2013).

ANCOVA was used to assess group-level differences in cognitive PC scores while adjusting for FA values from the arcuate fasciculus and corpus callosum subregions (splenium, genu, and body) as covariates. The dependent variable was cognitive PC score, and the independent variable was lateralisation group, tested under two comparisons: (1) typical, atypical, and strongly atypical groups and (2) cross-dominant and consistently lateralised groups. In addition to these group-level analyses, multiple regression was employed to examine within-group associations between FA and cognitive PCs. Separate regression models were fitted for each lateralisation group and for each tract to investigate potential tract-specific effects.

All analyses were conducted under two covariate conditions: (1) models including age and sex and (2) models additionally including handedness, as previous studies have suggested that handedness modulates the relationship between FA in language-related tracts and lateralisation (Andrulyte, Demirkan, et al., 2025).

## Results

### Demographic characteristics and language lateralisation

Participants were categorised based on task-based fMRI functional language lateralisation and patterns of crossed language dominance. For language lateralisation, participants were grouped into three categories: typical (n = 235), atypical (n = 35), and strongly atypical (n = 9). The gender distribution was similar across the groups (x^2^(2,N = 279) = 2.32, p = 0.31), and there were no significant differences in age (F(2,276) = 1.218, p =0.298). However, handedness scores differed significantly (F(2,276) = 8.52, p<0.001; Table 1).

Participants were also classified into consistent (n = 261) and crossed dominance (n = 18) groups based on frontal and temporal lateralisation patterns. Gender distribution did not differ significantly between these groups (x^2^(1,N = 279) = 0.10, p = 0.75). There were no significant differences in age (t(18.98) = −0.39, p = 0.701) or handedness scores (t(19.66) = 0.82, p = 0.422; Table 2).

### Principal components of cognitive scores

The PCA conducted on cognitive task scores (Table 3) identified three PCs that together explained 58.5% of the total variance in task performance (Figure 2A). PC1 accounted for 31.5% of the variance, with all task loadings being negative, indicating that this component was inversely associated with task scores. The tasks contributed similarly overall, with the most negative contributions from Raven’s matrices (-0.453), mental rotation (-0.403), and labyrinth navigation (-0.366). Based on the nature of these tasks, PC1 was interpreted as general cognitive function. For ease of interpretation, PC1 scores were inverted so that higher values correspond to better overall cognitive functioning. PC2, which explained 14.8% of the variance, loaded positively on labyrinth navigation (+0.409), mental rotation (+0.358), and Raven’s matrices (+0.169). This component was interpreted as reflecting visuospatial processing and memory. PC3, accounting for 12.2% of the variance, showed high positive loadings for arithmetical facts (+0.361) and complex mental calculations (+0.217). It was interpreted as representing mathematical ability.

**Figure 2.**
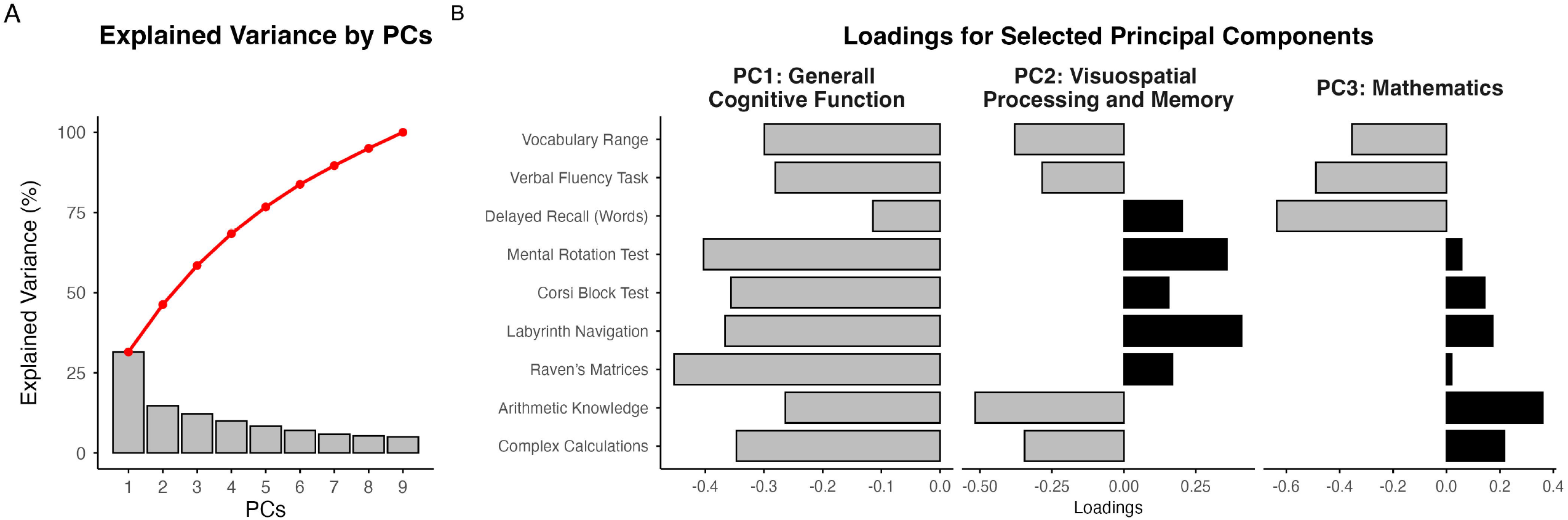
Principal component analysis (PCA) plots illustrating the explained variance and component loadings in the total sample. The PCA, conducted on cognitive test data, identified nine principal components, with the first three components collectively explaining 58.5% of the total variance. Panel (A) shows the proportion of variance explained by each component (bars) and the cumulative variance (red line) across all components. Panel (B) displays the loadings of individual cognitive tests on the first three components: PC1 (General Cognitive Function), PC2 (Visuospatial Processing and Memory), and PC3 (Mathematics). Bars to the left of the origin represent negative loadings, while those to the right represent positive loadings. Key cognitive tests contributing to each component are indicated, highlighting the cognitive tasks associated with each principal component.

### Typical and atypical language lateralisation

ANCOVA revealed no significant group-level differences in cognitive PC scores between typical, atypical, and strongly atypical lateralisation groups (pFDR > 0.05) (Supplementary Table 1). Multiple regression analyses showed a significant positive association between mean FA in the SCC and PC1 in individuals with strongly right-lateralised (i.e., strongly atypical) language dominance (R^2^ = 0.96, p = 0.002, pFDR = 0.042) (Figure 3A). Similarly, a positive relationship was observed between mean FA in the GCC and PC2 in this group (R^2^ = 0.92, p = 0.0009, pFDR = 0.042) (Figure 3C). Notably, these associations were not found in individuals with typical or atypical language lateralisation (pFDR > 0.05) (Supplementary Figure 1A and B). Interestingly, both the SCC-PC1 and GCC-PC2 associations lost significance (pFDR = 0.013) when handedness was included as a covariate (Figure 3B and D). No other significant associations were found after FDR correction (Supplementary Table 2).

**Figure 3.**
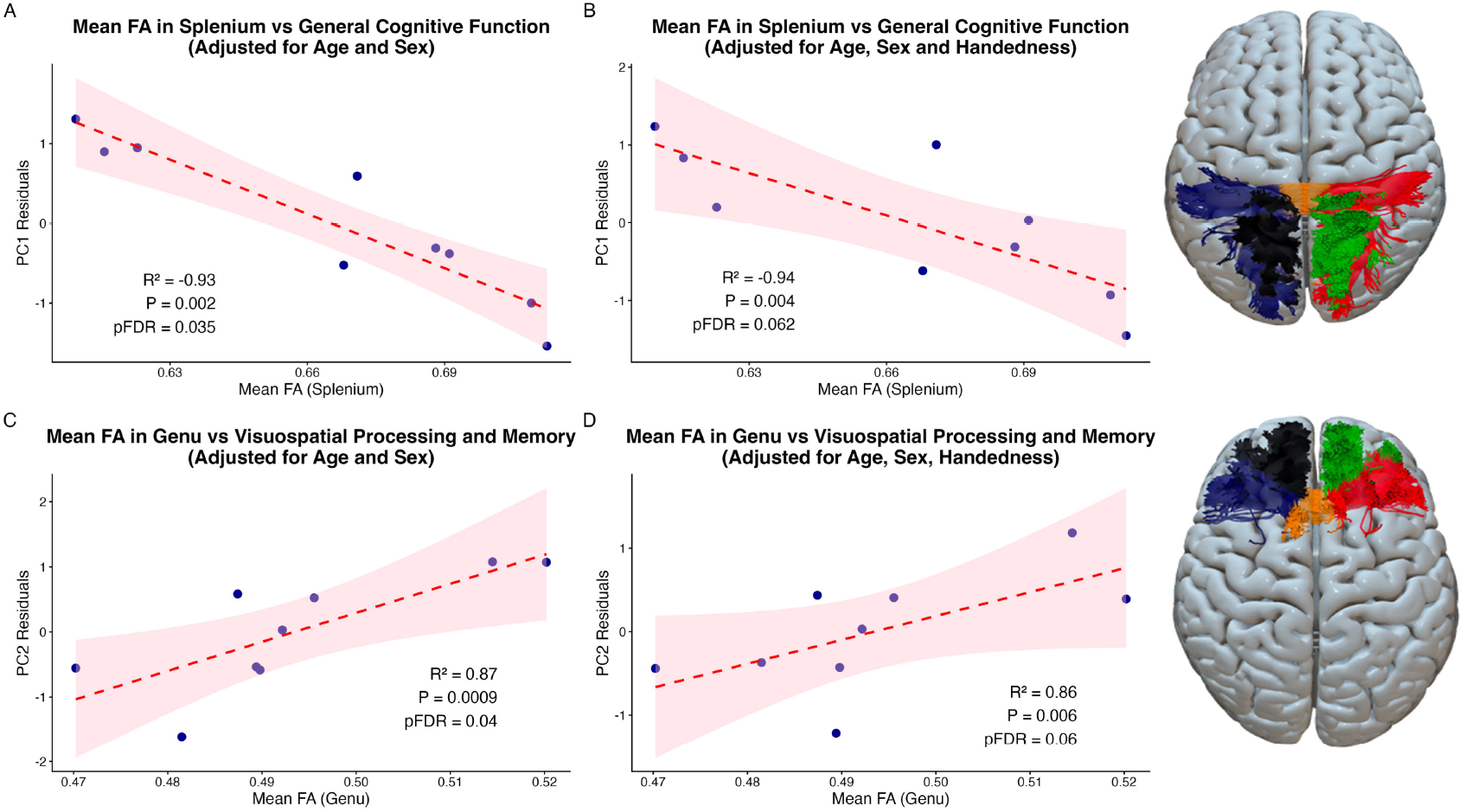
Associations between mean FA of the corpus callosum splenium and genu with cognitive scores in strongly atypical (right-lateralised) individuals. The regression plots illustrate the relationships between mean FA values of the corpus callosum (x-axis) and principal component scores (y-axis), adjusted for different covariates. Panels (A) and (B) examine the association between the mean FA of the corpus callosum splenium and general cognitive function (PC1), while panels (C) and (D) focus on the relationship between the mean FA of the corpus callosum genu and visuospatial processing and memory (PC2). In panel (A), there is a significant negative association between mean FA in the splenium and executive function PC1 scores after adjusting for age and sex (R^2^ = 0.96, pFDR = 0.042). However, in panel (B), when handedness is included as an additional covariate, this relationship does not survive correction for multiple comparisons (R^2^ = 0.97, pFDR = 0.129). Panel (C) shows a significant positive association between mean FA in the genu and visuospatial processing and memory PC2 scores, corrected for age and sex (R^2^ = 0.92, pFDR = 0.042). Again, this association becomes non-significant when handedness is added as a covariate in panel (D) (R^2^ = 0.93, pFDR = 0.129). The red dashed lines represent the regression fits, while the pink shaded areas indicate 95% confidence intervals. The reconstructed corpus callosum bundles on the right show the regions of interest: the splenium in the top image and the genu in the bottom image. For a detailed comparison with individuals exhibiting left or bilateral language representation, refer to Supplementary Figure 1.

**Figure 4.**
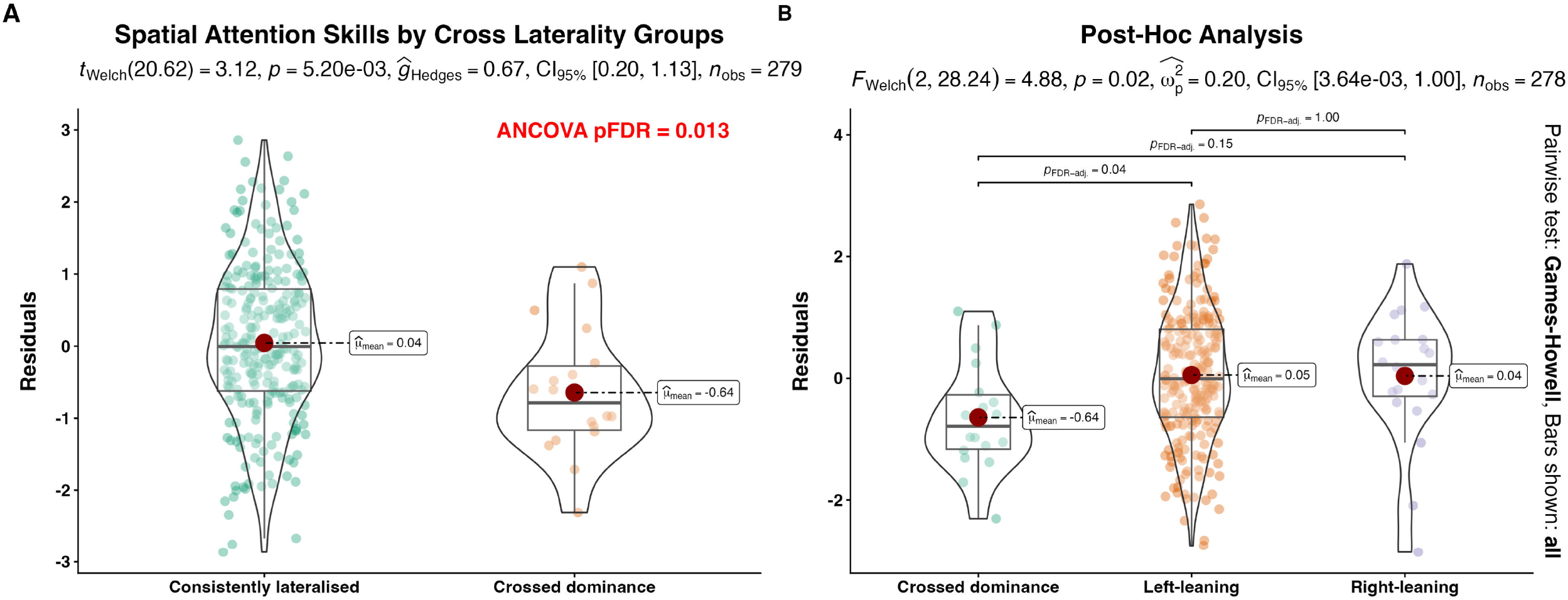
Association between consistency in language lateralisation and spatial attention cognitive scores. (A) ANCOVA, adjusted for handedness, sex, and age, found a significant main effect of cross-laterality group on visuospatial processing and memory scores (PC2) (F = 8.45, pFDR = 0.013). Violin plots show the residuals of visuospatial processing and memory scores (PC2) for consistently lateralised and cross-dominant individuals (adjusted for age, sex, and handedness). (B) Post-hoc analyses revealed that consistently left-lateralised individuals had significantly higher spatial attention scores compared to cross-dominant individuals (pFDR = 0.02), while no significant difference was observed between consistently right-lateralised and cross-dominant groups (pFDR = 0.22). Individual participant values are shown as points, with large red points indicating group means. Games-Howell test was used for post-hoc pairwise comparisons and is indicated by error bars.

### Crossed language dominance

ANCOVA revealed a significant main effect of cross-laterality group on PC2 (visuospatial processing and memory) (F(2, 278) = 8.45, pFDR = 0.02), with crossed-dominant individuals showing significantly lower PC2 scores compared to consistently lateralised individuals (t(279) = 3.4, p = 0.003). These results remained significant regardless of the inclusion of handedness as a covariate (Supplementary Table 3). No other significant ANCOVA results were observed, including interaction effects between FA and laterality (all pFDR > 0.05).

To better understand which laterality group was driving this effect, post-hoc analyses were conducted. A one-way ANOVA confirmed a significant main effect of laterality (F(2, 277) = 8.51, p = 0.004). Games-Howell post-hoc tests revealed that crossed-dominant individuals had significantly lower PC2 scores than consistently left-lateralised individuals (pFDR = 0.02). No significant differences were observed between crossed-dominant and consistently right-lateralised individuals (pFDR = 0.22), nor between consistently left- and consistently right-lateralised groups (pFDR = 0.94).

Unlike the group-level differences identified in ANCOVA, multiple regression analyses did not reveal significant associations between FA and cognitive PC scores in either crossed-dominant or consistently lateralised individuals (pFDR > 0.05) (Supplementary Table 2).

## Discussion

This study is the first to investigate the relationship between FA in the arcuate fasciculus and corpus callosum and cognitive performance across distinct language lateralisation groups. Our findings reveal three key results. First, while cognitive performance did not differ between typical, atypical, and strongly atypical groups, corpus callosum FA was positively associated with general cognition and visuospatial processing in strongly right-lateralised individuals. This relationship was not observed in those with bilateral or leftward lateralisation, and notably, it was no longer significant when handedness was included as a covariate. Second, individuals with consistent language dominance exhibited higher visuospatial cognitive scores compared to those with crossed language dominance, an effect that emerged independently of FA. Importantly, this cognitive difference was driven by individuals with consistent leftward lateralisation, whereas no significant differences were observed between crossed dominance and consistent rightward lateralisation. Finally, FA in the left or right arcuate fasciculus did not moderate any of these associations.

### Cognitive implications of atypical lateralisation

Three primary mechanisms have been proposed to explain how hemispheric asymmetries enhance cognitive efficiency: (1) preferential engagement of one hemisphere increases its functional efficiency; (2) minimising interference from the non-dominant hemisphere improves reaction times; and (3) asymmetry enables parallel processing of complementary information across hemispheres, enhancing multitasking capacity (Güntürkün et al., 2020). If left-lateralisation optimises cognitive processing within a single hemisphere, it follows that left-lateralised individuals require less interhemispheric connectivity, explaining the absence of significant corpus callosum FA-cognition associations in this group. Their intrahemispheric specialisation may already provide sufficient efficiency for general cognition and visuospatial tasks.

Although atypical language lateralisation, particularly rightward dominance, has been associated with neurodevelopmental disorders and cognitive deficits (Villar-Rodríguez, Cano-Melle, et al., 2024), we did not find such associations when examining cognitive performance in our study. This aligns with previous research on healthy individuals, suggesting that atypical patterns represent alternative neural strategies rather than inherent dysfunctions. Therefore, they should not be considered maladaptive but rather a variant of neural organisation with potential compensatory advantages (Illingworth & Bishop, 2009; Knecht, 2001; Whitehouse & Bishop, 2008). Evolutionary perspectives suggest that population-level asymmetries arise due to selective pressures. While left-hemisphere dominance is predominant, rightward language dominance may confer alternative benefits. Although far from the complexities of the human brain, evidence from lateralised species such as fish supports this idea: collective directional movement reduces predation risk, but individuals with reversed asymmetry may gain an advantage by behaving unpredictably and avoiding anticipated predator attacks (Vallortigara & Rogers, 2005). Applying this framework to humans, the predominance of left-hemisphere language dominance may reflect evolutionary pressures favouring intrahemispheric specialisation, whereas rightward dominance could represent an alternative strategy, potentially supporting greater cognitive flexibility through increased interhemispheric integration

### Corpus callosum

The corpus callosum is the principal interhemispheric communication pathway, facilitating information transfer between homologous cortical regions (Ocklenburg & Guo, 2024). Structural and functional MRI studies suggest that right-lateralised individuals exhibit stronger interhemispheric connectivity compared to typically lateralised individuals (Andrulyte, Zago, et al., 2025; Labache et al., 2020), aligning with evidence that the left hemisphere maintains denser intrahemispheric connections, while the right hemisphere relies more on interhemispheric integration (Gotts et al., 2013). The observed positive association between corpus callosum FA and cognitive performance in right-lateralised individuals suggests an adaptive mechanism supporting general cognitive and visuospatial functions.

Our results further demonstrate regional specificity within the corpus callosum. Increased FA in the splenium was associated with better overall cognitive function, while greater FA in the genu correlated with higher visuospatial scores, findings consistent with previous studies (Bhadelia et al., 2009; Fryer et al., 2008). The splenium, which connects occipital, parietal, and posterior temporal regions, facilitates interhemispheric communication crucial for visual and linguistic processing, including picture naming (Laporta-Hoyos et al., 2023), syntactic processing (Midrigan-Ciochina et al., 2024), and expressive vocabulary (Fryer et al., 2008). The genu, connecting frontal cortices, is implicated in higher-order cognitive functions such as attentional control and visuospatial coordination (Bhadelia et al., 2009; McFarland & Haber, 2002). Interestingly, Zhu et al. (2025) demonstrated that anterior corpus callosum structure is linked to spatial attention lateralisation, although their findings primarily involved the rostrum rather than the genu.

From a neurodevelopmental perspective, hemispheric lateralisation typically increases with age, as the brain refines functional specialisations to optimise cognitive efficiency (Olulade et al., 2020; Ringo et al., 1994). Some theories propose that deviations from this trajectory arise from developmental instability, where genetic or environmental factors disrupt systematic lateralisation patterns (Yeo et al., 1997; Yeo & Gangestad, 2015). However, our results suggest a more structured relationship: rather than reflecting random variability, increased callosal FA in right-lateralised individuals may compensate for functional demands. Given that both general cognition and visuospatial cognitive performance correlated with corpus callosum FA in this group, one interpretation is that right-lateralised individuals experience a higher functional load within the right hemisphere (Hervé et al., 2013), necessitating enhanced interhemispheric transfer to mitigate potential resource competition. This aligns with the cognitive crowding hypothesis, which posits that lateralising multiple cognitive functions to the same hemisphere creates competition for neural resources (Groen et al., 2012). Supporting this, studies have found that right-lateralised individuals with both language and visuospatial functions lateralised to the right hemisphere do not exhibit deficits in executive function or spatial cognition (Flöel et al., 2001) and are associated with increased FA in the corpus callosum (Häberling et al., 2011). However, as our study did not look at the visuospatial lateralisation, these interpretations remain speculative.

While atypical language dominance—whether rightward or bilateral—was not associated with reduced cognitive performance, individuals with crossed dominance exhibited significantly lower visuospatial attention scores compared to consistently lateralised individuals. Unlike right-lateralised individuals, who appear to compensate for their atypical language organisation through enhanced interhemispheric communication, crossed-dominant individuals may experience greater demands for neural coordination between hemispheres without sufficient structural support (Yeo et al., 1997). However, if competition for neural resources were the primary factor, a similar cognitive disadvantage would be expected in right-lateralised individuals, yet no such reduction in visuospatial performance was observed. This suggests that the observed effect is not simply a result of cognitive crowding within a single hemisphere but rather a consequence of disrupted neural integration within the language system itself, also known as the language laterality profile hypothesis, which suggests that the risk of language difficulties (e.g., dyslexia) increases when different linguistic functions are predominantly mediated by opposite hemispheres rather than being lateralised in a coordinated manner (Illingworth & Bishop, 2009).

Although our study did not examine dyslexia, our findings indicate that a similar lack of consistency in lateralisation across language domains may introduce inefficiencies in cognitive processing. Even though a single language production task was employed, it engages multiple interdependent components (Bradshaw, Thompson, et al., 2017), each typically associated with distinct neural regions (Hodgson et al., 2021). Thus, rather than a generalised effect of atypical lateralisation, our findings suggest that specific patterns of hemispheric specialisation—such as crossed dominance—may contribute to cognitive inefficiencies. Future research should investigate whether crossed-dominant individuals exhibit altered functional connectivity between frontal and temporal regions and whether similar disadvantages extend to other cognitive domains beyond visuospatial attention.

### Handedness

Handedness and language lateralisation are consistently associated, yet their precise neurodevelopmental relationship remains uncertain (Ocklenburg et al., 2014). Handedness is often used as a proxy for hemispheric dominance, with approximately 95% of right-handers and 70–85% of left-handers exhibiting left-hemispheric language dominance (Knecht, Dräger, et al., 2000), yet it explains only a small proportion of variance in lateralisation (r^2^ ≈ 0.13; Badzakova-Trajkov et al., 2010). Beyond language, handedness is also linked to white matter organisation, with larger corpus callosum associated with lower handedness scores (although with a small effect size) (Josse et al., 2008; Luders et al., 2010). In our study, handedness was included as a covariate alongside FA to examine its influence on cognitive performance within each laterality group, and despite not being the primary focus, its inclusion altered the strength of FA-cognition associations. For instance, in the splenium, the relationship between FA and general cognitive function weakened from p = 0.002 to p = 0.004, with the FDR-adjusted value shifting from pFDR = 0.042 to pFDR = 0.129, while in the genu, the FA-visuospatial processing association saw a similar attenuation (p = 0.0009 to p = 0.006; pFDR = 0.042 to pFDR = 0.129). While these raw p-value changes may appear subtle, their impact on FDR correction suggests that the variance removed by adjusting for handedness was not merely statistical noise but may have overlapped with meaningful biological variability underlying the relationships between corpus callosum structure, language lateralisation, and cognition.

One plausible explanation is Lord’s paradox, where statistical adjustment for a variable that shares neurodevelopmental underpinnings with both the predictor and outcome can lead to an overcorrection, inadvertently masking genuine effects (Tu et al., 2008). If handedness, corpus callosum FA, and right-hemispheric lateralisation for language are interdependent, controlling for handedness may have removed meaningful variance rather than providing a clearer estimate of FA-cognition associations. However, an alternative interpretation is that handedness plays an independent role in cognitive function, rather than merely acting as a confounder. Right-handers have been reported to exhibit small but consistent cognitive advantages in visuospatial processing tasks (Somers et al., 2015), raising the possibility that handedness contributes directly to interhemispheric communication dynamics. Under this view, corpus callosum FA might differentially support cognition depending on handedness, rather than simply reflecting a statistical artefact. The observed effects in our study, though likely driven by Lord’s paradox given their small magnitude, could nonetheless reflect a genuine interaction between handedness and CC function.

### Methodological considerations

A key limitation of this study is the small sample size (n = 9) for the strongly right-lateralised group. While the exceptionally high R^2^ (>0.9) observed between corpus callosum FA and general cognition and visuospatial processing does not invalidate our findings, it should be interpreted with caution. Small sample sizes, particularly when multiple covariates are included, can increase the risk of overfitting and inflated effect sizes (Button et al., 2013). Additionally, we did not assess visuospatial lateralisation, limiting direct evaluation of the cognitive crowding hypothesis. Future studies should incorporate functional laterality measures alongside structural imaging to clarify these relationships.

Another important consideration concerns the limitations of the diffusion MRI acquisition used in the BIL&GIN dataset. Although valuable, the single-shell, low angular resolution DTI protocol employed here constrains the interpretability of microstructural features, particularly in regions with crossing fibres. Future studies would benefit from adopting high angular resolution diffusion imaging (HARDI) with multi-shell acquisitions, enabling advanced models such as constrained spherical deconvolution (CSD) and fixel-based analysis (Jeurissen et al., 2014). These approaches allow for improved resolution of fibre populations and more specific characterisation of white matter architecture. Moreover, the integration of complementary techniques such as myelin-sensitive imaging (e.g., magnetisation transfer, quantitative T1 mapping) could offer further insight into the biological underpinnings of structural asymmetries (Trampel et al., 2019). Despite the aforementioned limitations, our findings offer novel insights into the associations between white matter FA, atypical lateralisation, and cognition. Larger, well-powered studies employing advanced structural and functional imaging protocols are needed to validate these associations and to elucidate whether increased corpus callosum FA in right-lateralised individuals reflects compensatory adaptation or an alternative neurodevelopmental pathway.

## Acknowledgements

We would like to thank Dr Emmanuel Mellet for his valuable contributions to this study. His input and support throughout the research process have been greatly appreciated. We also acknowledge the financial support of the Biotechnology and Biological Sciences Research Council (BBSRC).

## Funding

Ieva Andrulyte – BBSRC DTP training grant BB/T008695/1

## CREDIT authorship contribution statement

Ieva Andrulyte – conceptualisation, data analysis, funding acquisition, writing – original draft

Laure Zago – data collection, writing – review & editing Gael Jobard – data collection, methodology

Herve Lemaitre – data collection

Francois Rheault – methodology, writing – review & editing Marc Joliot – data collection, methodology, writing – review

Laurent Petit – data collection, methodology, writing – review & editing, supervision

Simon S Keller – conceptualisation, writing – review & editing, supervision

## Competing Interests

Authors declare no competing interests

## Data availability statement

All analysis scripts used in the present study are openly available on GitHub: https://github.com/andrulyte/language-laterality-cognition.

## Notes

### Competing Interest Statement

The authors have declared no competing interest.

### Summary of Updates

To update the quality of the figures and include GitHub information.

