## Supplementary figures and images for "Language laterality and cognitive skills: does anatomy matter?"

### Supplementary Figure 1

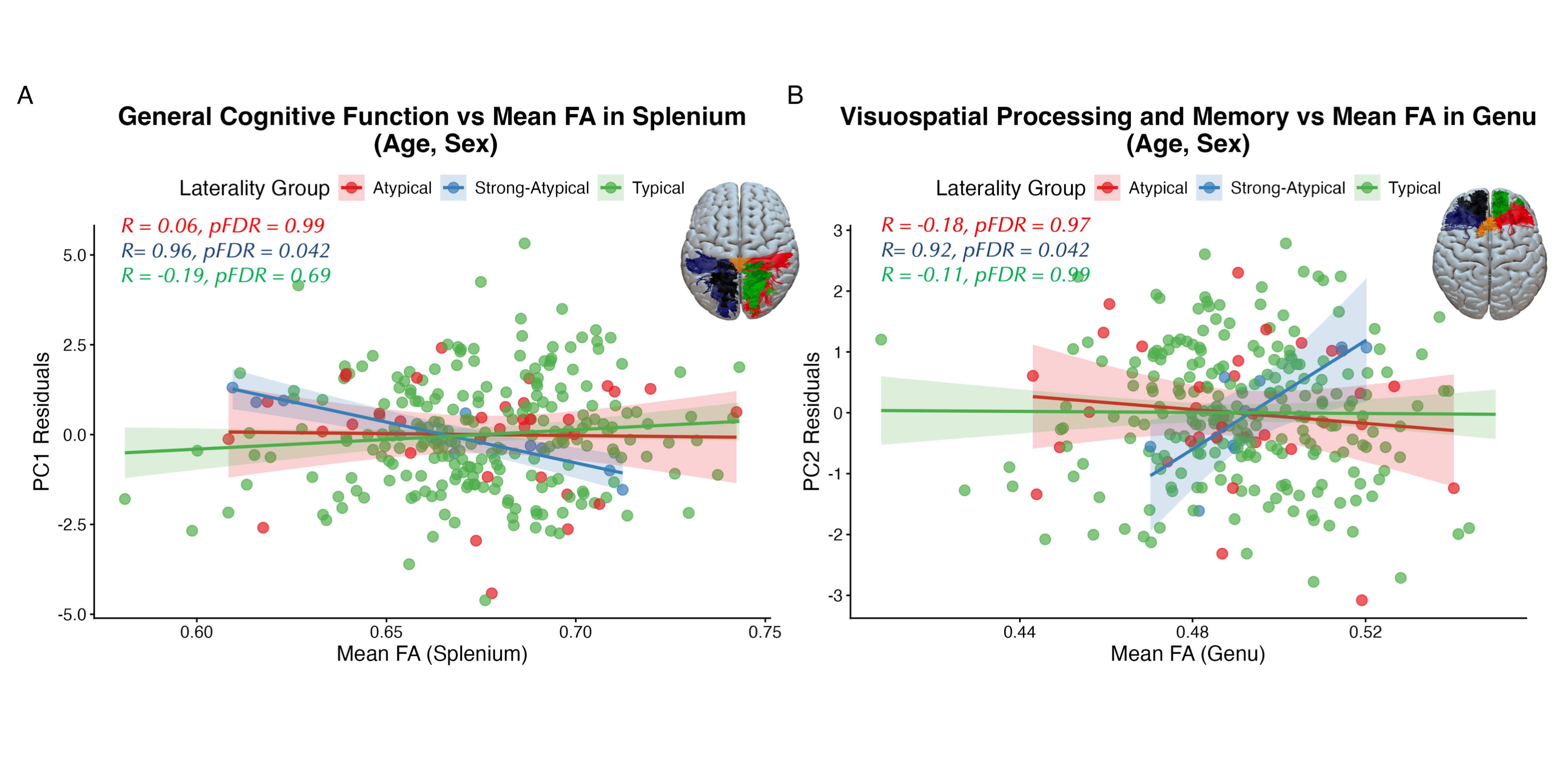
