## Supplementary Figure Legend for "Language laterality and cognitive skills: does anatomy matter?"

**Supplementary Figure 1. Associations between cognitive performance and mean FA in the corpus callosum splenium and genu across language lateralisation groups.** (A) Regression plots illustrating the relationship between residual executive function principal component (PC1) scores (y-axis) and mean FA in the splenium of the corpus callosum (x-axis), adjusted for age and sex. (B) Regression plots depicting the association between residual spatial attention principal component (PC2) scores (y-axis) and mean FA in the genu of the corpus callosum (x-axis), also adjusted for age and sex. The regression lines and shaded 95% confidence intervals represent the associations stratified by language lateralisation groups: atypical (red; bilateral to mildly right lateralised), strongly atypical (blue; moderately to strongly right lateralised), and typical (green; left lateralised). Only the strongly atypical group exhibited a significant relationship between cognitive scores and mean FA in both the splenium and genu, whereas no significant associations were observed in the atypical and typical groups. Brain schematic inserts display reconstructions of the splenium (A) and genu (B) regions of the corpus callosum for reference. R squared values and corresponding false discovery rate-adjusted p-values (pFDR) are provided for each group.
